# The crystal structure of *Vibrio cholerae* (6-4) photolyase reveals interactions with cofactors and a DNA binding region

**DOI:** 10.1101/2022.09.10.507404

**Authors:** Baris Cakilkaya, Ibrahim Halil Kavakli, Hasan DeMirci

## Abstract

Photolyases (PLs) reverse UV-induced DNA damage using blue light as an energy source. Of these photolyases, (6-4) PLs repair (6-4)-lesioned photoproducts. We recently identified a gene from *Vibrio cholerae (Vc*) encoding a (6-4) PL, but structural characterization is needed to elucidate specific interactions with the chromophore cofactors. Here, we determined the crystal structure of *Vc*(6-4) PL at 2.5 Å resolution. Our high-resolution structure revealed that the two well-known cofactors, flavin adenine dinucleotide (FAD) and the photoantenna 6,7-dimethyl 8-ribityl-lumazin (DMRL), stably interact with an α-helical and an α/β domain, respectively. Additionally, the structure has a third cofactor with distinct electron clouds corresponding to a [4Fe-4S] cluster. Moreover, we identified that Asp106 makes a hydrogen bond with the water and DMRL, which indicates further stabilization of the photoantenna DMRL within *Vc*(6-4) PL. Further analysis of the *Vc*(6-4) PL structure revealed a possible region responsible for DNA binding. The region located between residues 478-484 may bind the lesioned DNA, with Arg483 potentially forming a salt bridge with DNA to stabilize further the interaction of *Vc*(6-4) PL with its substrate. Our comparative analysis revealed that the DNA lesion could not bind to the *Vc*(6-4) PL in a similar fashion to the *Drosophila melanogaster Dm*(6-4) PL without a significant conformational change of the protein. The 23^rd^ helix of the bacterial (6-4) PLs seems to have remarkable plasticity, and conformational changes facilitate DNA binding. In conclusion, our structure provides further insight into DNA repair by a (6-4) PL containing three cofactors.

## Introduction

The cryptochrome/photolyase family (CPF) is a large family of flavoproteins that absorb near UV–visible light (300–500 nm range) and has diverse functions depending on the type of organism (1, 2). Cryptochromes (CRYs) are the circadian photoreceptors in *Drosophila* and plants while they are acting as transcriptional repressor of the circadian clock in mammals (2–4). Photolyases (PLs) are enzymes that repair UV-induced DNA damage (5–7). Although phylogenetic analyses revealed that there are different classes of PLs, there are three different types of PLs based on their repair activities: Cyclobutane pyrimidine dimer (CPD) photolyases (PLs), (6–4) photolyases, and *Drosophila, Arabidopsis, Synechocystis*, human (DASH)-type cryptochrome (CRY-DASH) (8–10). CPD PLs repair mainly cyclobutane pyrimidine dimers (Pyr<>Pyr) while (6-4) PLs repair pyrimidine-pyrimidine photoproducts (Pyr [6–4] Pyr) using blue light (350–500 nm) as the energy source (1, 6). CRY-DASHs have quite diverse physiological functions including single-strand DNA photolyase activity (11). PLs are monomeric proteins with molecular masses of 50–61 kDa and possess two chromophores (12). Flavin Adenine Dinucleotide (FAD) is an essential cofactor for the catalytic activity of PLs and acts as a catalytic cofactor in all PLs (5, 12). The second chromophore, which acts as photoantenna, varies depending on the organisms. Following photoantenna are being found: methenyltetrahydrofolate (MTHF), 8- hydroxy-7,8-didemethyl-5-deazariboflavin (8-HDF), FAD, flavin mononucleotide (FMN), or 6,7- dimethyl 8-ribityl-lumazin (DMRL) depending on the type of photolyase (13, 14). The (6–4) photolyases had previously been thought to exist only in eukaryotes. However, studies with different bacteria revealed the presence of (6-4) PLs (15–17). These bacterial (6-4) PLs are unique having [4Fe-4S] in addition to DMRL and FAD cofactors (15, 17).

Crystal structures of various PLs have been determined by X-ray crystallography. First available crystal structure is the *Escherichia coli* CPD PL (18). Subsequent structures have been obtained from different organismal PLs including *Anacystis nidulans* (19), *Thermus thermophilus* (20), (6–4) photolyases of *Arabidopsis thaliana* (21) and *Agrobacterium tumefaciens* (13). Comparison of these crystal structures of various homologous PLs revealed valuable information about their structural features and enzyme mechanisms. PLs are made up of two well-defined domains which are a α/β and an α-helical domains. The α-helical domain is associated with FAD while α/β domain is the binding site for the second chromophore (12). These two domains are connected by a long flexible loop.

*V. cholerae* O1 bivar Tor str. N16961 has a CPD photolyase that repairs T<>T dimers and two CRY-DASHs that repair T<>T dimers in single-stranded DNA (11, 22). Our previous study with a gene (*Vca0809*) in *V. cholerae* highly expressed upon exposure of the organism to blue light (23). Our further characterization of this gene revealed that it encodes (6-4) PL with FAD and DMRL and consists of the [4Fe-4S] domain (15). Here, we determined the crystal structure of *Vc*(6-4) PL to 2.5 Å resolution. Our crystal structure comparison revealed a putative DNA binding region of *Vc*(6-4) PL.

## Results

### Crystal structure of *Vibrio cholerae* (6-4) photolyase

We recently identified a (6-4) PL from *Vibrio cholerae* O1 bivar Tor str. N16961(*Vc*) which possesses catalytic cofactor FAD, an antenna chromophore DMRL, and additional cofactor [4Fe-4S] cluster (15). We determined the crystal structure of *Vc*(6-4) PL at 2.5 Å resolution at cryogenic temperature at the Turkish Light Source known as *“Turkish DeLighf* (24, 25). The structure contains all the cofactors with well-defined electron densities and contains a total of 24 α helixes and 5 β sheets (**Figure 1A**). It consists of two domains: an α-helical domain that is associated with FAD and α/β domain which interacts with DMRL cofactor. The sequence alignment of *Vc*(6-4) PL with structurally available bacterial (6-4) PL with 38.4% sequence similarity with *Agrobacterium tumefaciens (At)(6-4*) PL and 43.1% sequence similarity with *Rhodobacter sphaeroides* (*Rs*)(6-4) PL (**Figure 1B**).

**Figure 1:**
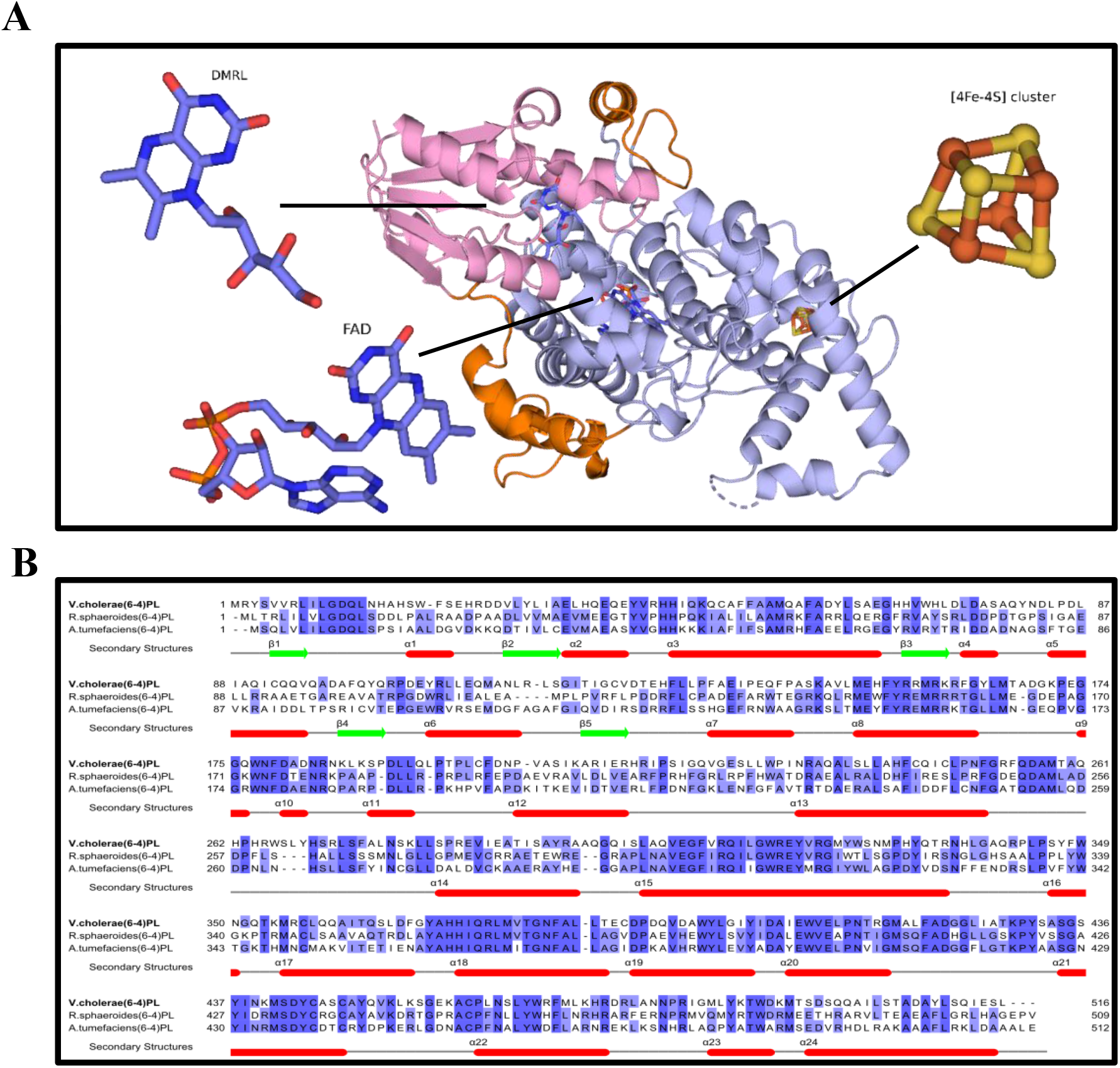
Crystal structure and sequence comparison of *Vc*(6-4) PL. (A) Overview of *Vc*(6-4) PL structure, pink is α/β-domain, light blue is catalytic domain, and orange is the inter-domain linker region. Cofactor places are indicated by the black line. (B) Sequence alignment of bacterial (6-4) photolyases with available crystal structures and secondary structures of *Vc*(6-4) PL are shown. Red lines are α helixes and green lines are β sheets. Percentage identity coloring is used in *JALVIEW* software.

### DMRL Binding domain

DMRL is a photoantenna chromophore and has higher efficiency to absorb light than FAD for increasing the catalytic efficiency (13). In our crystal structure, there is a very well-defined electron density of the DMRL and interacting amino acid residues (**Figure 2A**). Aromatic ring of DMRL interacts with nitrogen atoms of Gln11, Leu35, Gln39, and oxygen atom of Ala33 (**Figure 2B**). Additionally, the aromatic ring of DMRL interacts with Trp488, Ala33 and Ile9. The ribityl group of the DMRL forms hydrogen bonds with oxygen atoms of the Glu38, Tyr41, Asp12 and with amino group of Asp10. When we compare the interaction partners of DMRL in *Vc*(6-4) PL with other known bacterial (6-4) PLs, almost all the interactions are well-conserved with a few exceptions. First, Cys32 carboxyl group of *At*(6-4) PL, interacts with aromatic ring in DMRL (PDB ID: 4DJA) while in both *Vc* and *Rs*(6-4) PLs DMRL interact with main chain carbonyl group of Ala33. We identified a critical water molecule (W22) that interacts with Asp12, Gln13, Asp106, and DMRL in *Vc*(6-4) PL (**Figure 2C**). Analyses of other bacterial (6-4) PLs have shown similar conserved motifs (**Figure 2C**). Such motifs increase the overall stability of the DMRL within the enzyme. However, the W22 molecule interacts with Asp106 of the *Vc*(6-4) PL with a distance of 2.8 Å while the same interaction occurs with Gly105 of other bacterial (6-4) PLs with a distance of over 3.3 Å. These differences might increase the affinity of DMRL with *Vc*(6-4) PL.

**Figure 2:**
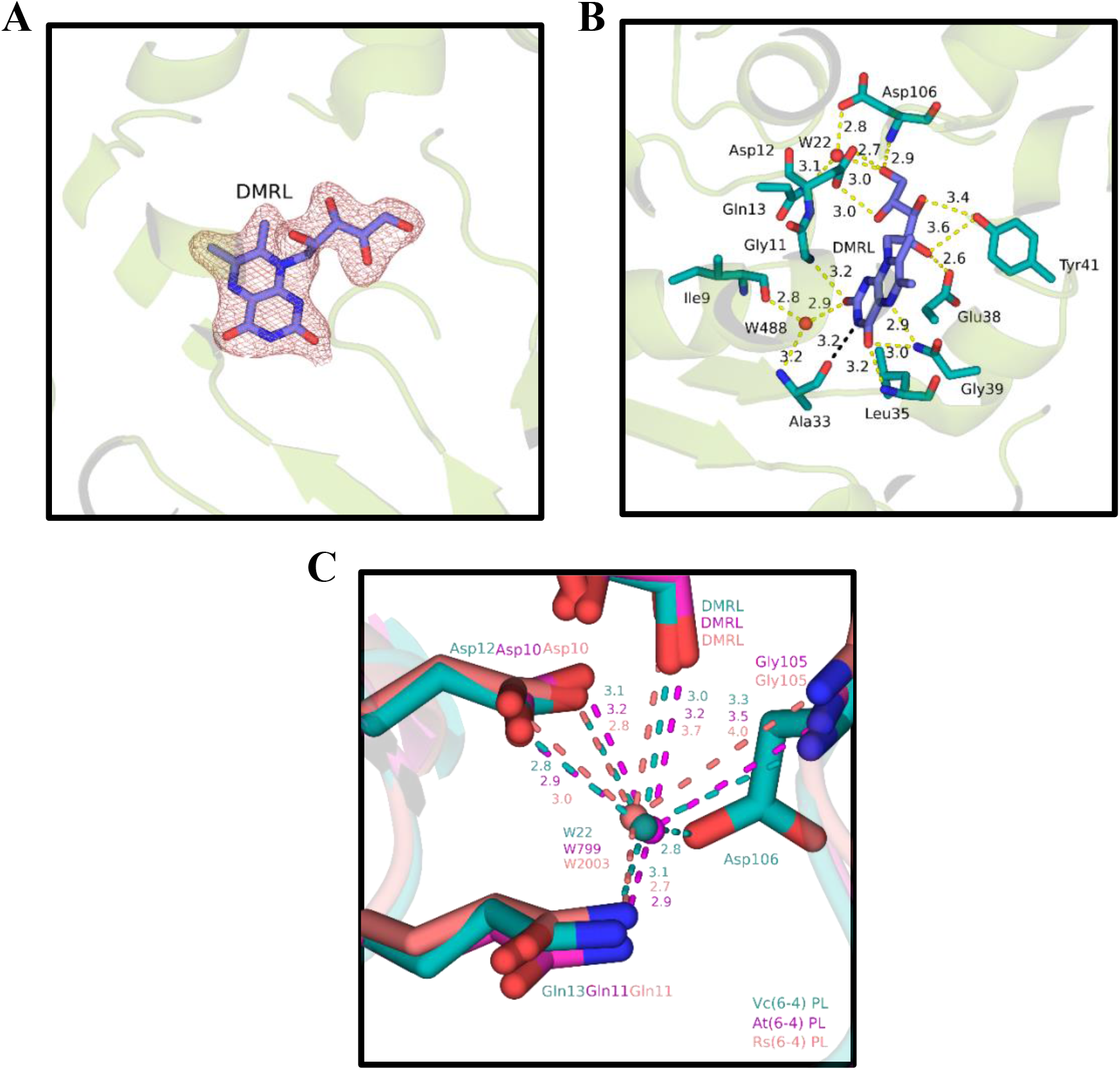
DMRL interactions and comparisons with other (6-4) photolyases. Every distance is in units of Armstrong (Å). (A) Electron cloud of DMRL at σA-weighted 2 Fo-Fc electron density map contoured at 1.0σ in deepsalmon mesh and *Vc*(6-4) PL cartoon in splitpea color. (B) Possible interaction partners of DMRL. Yellow dashes represent hydrogen bondings, black dashes indicate Van der Waals interactions, and red dots are water molecules. (C) Additional water molecules that might stabilize the DMRL interaction. Deepteal, magenta, and deepsalmon colors indicate *Vc*(6- 4) PL, *At*(6-4) PL (PDB ID: 4DJA) and *Rs*(6-4) PL (PDB ID: 3ZXS) respectively. Their superimposed structures have RMSD: 0.834 Å (*At*(6-4) PL), and RMSD: 0.876 Å (*Rs*(6-4) PL). Each dashed line represents hydrogen bonding of species with corresponding color.

### FAD binding domain

FAD is located in the α-helical domain with U shaped conformation (**Figure 3A**). The N5 and O4 of FAD make hydrogen bonds with W7 and additional hydrogen bonds with carbonyl group of Tyr398 and NH1 atom of Arg376 residue (**Figure 3B**). As previously suggested for other PLs (17) these critical interactions may also serve a stabilization function *Vc*(6-4) PL. Unlike *At*(6- 4) PL or *Rs*(6-4) PL, *Vc*(6-4) PL Glu410 amino acid faces away from FAD and interacts with Asp395 rather than an His365 residue (**Figure 3B**). This results in a much-relaxed interaction between the residues due to hydrogen bonding formation rather than a salt bridge formation in *Vc*(6-4) PL.

**Figure 3:**
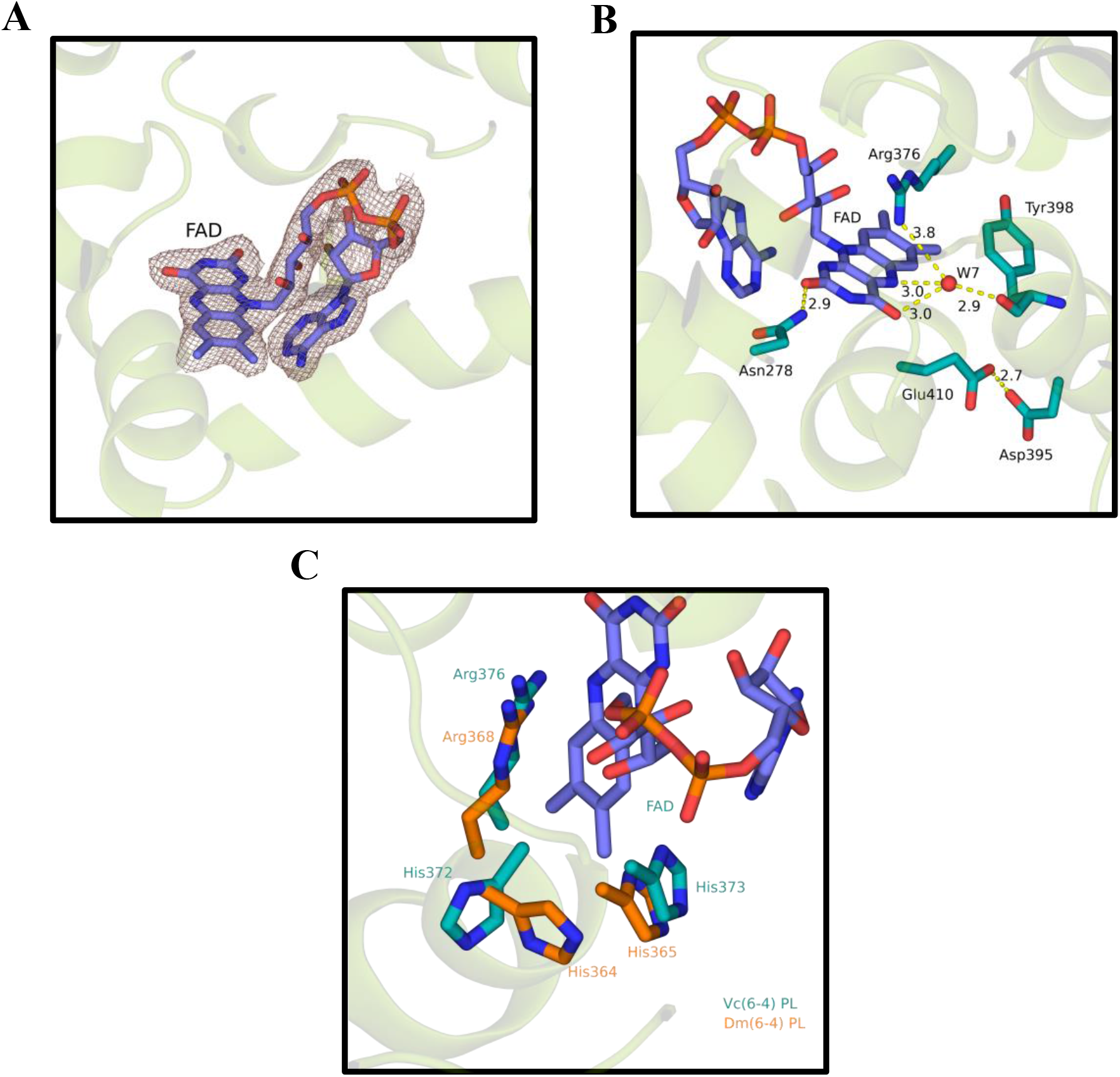
FAD interactions and comparisons with other (6-4) photolyases. Every distance is in units of Angstrom (Å). (A) Electron cloud of FAD at σA-weighted 2 Fo-Fc electron density map contoured at 1.0σ in deepsalmon mesh, and *Vc*(6-4) PL cartoon in splitpea color. (B) Possible interaction partners of FAD. Yellow dashes represent hydrogen bondings, red dots are water molecules. (C) Superimposed structure of *Vc*(6-4) PL and *Dm*(6-4) PL (PDB ID: 7AYV) at RMSD: 4.350 Å. Teal color represents *Vc*(6-4) PL and tv_orange color represents *Dm*(6-4) PL.

Although the reaction mechanism of the CPD photolyases is elucidated, (6-4) PLs not yet fully characterized. There are different mechanisms that are proposed for the (6-4) PL mechanism during DNA repair (7, 14, 26 & 27). In fact, mutagenesis studies with *Dm*(6-4) PL indicated that replacement of His365 to Asn365 result in loss of its DNA repair activity while mutagenesis of His369 into Met369 result in highly reduced activity of *Dm*(6-4) PL (28). However, mutagenesis studies with corresponding amino acid residues in *Xenopus* (6-4) PL showed complete loss of activity (29). Femtosecond spectroscopy and site-directed mutagenesis suggest initial electron transfer from excited flavin induces transfer of a proton from a histidine on the active site of the enzyme to the (6-4) photoproduct (7, 30).

Catalytically active His373 (corresponds to His365 of the *Dm*(6-4) PL) is conserved in *Vc*(6-4) PL (**Figure 3C**). In addition, there are other conserved amino acid residues in the motif. For instance, His372 and Arg376 (corresponds to His364 and Arg368 of the *Dm*(6-4) PL) are conserved in *Vc*(6-4) PL (**Figure 3C**). However, the conformation of the *Vc*(6-4) PL His372 is significantly different from the *Dm*(6-4) PL His364, which may suggest an alternate function of this residue.

### [4Fe-4S] cluster

The high-resolution Vc(6-4) PL structure contains an iron-sulfur cluster with an unknown function with a well-defined electron density (**Figure 4A**). Its interactions with the surrounding cysteine amino acids summarized in **Figure 4B**. Cys357, Cys445, Cys448, Cys461 sulfide atoms make bonds with Fe atoms in the cluster with each bond at 2.3 Å distance. The function of the [4Fe-4S] cluster is currently unknown, however, its distance to the FAD is 17 Å, which indicates it does not have a catalytic role in the DNA repair due to its large distance to the active site region.

**Figure 4:**
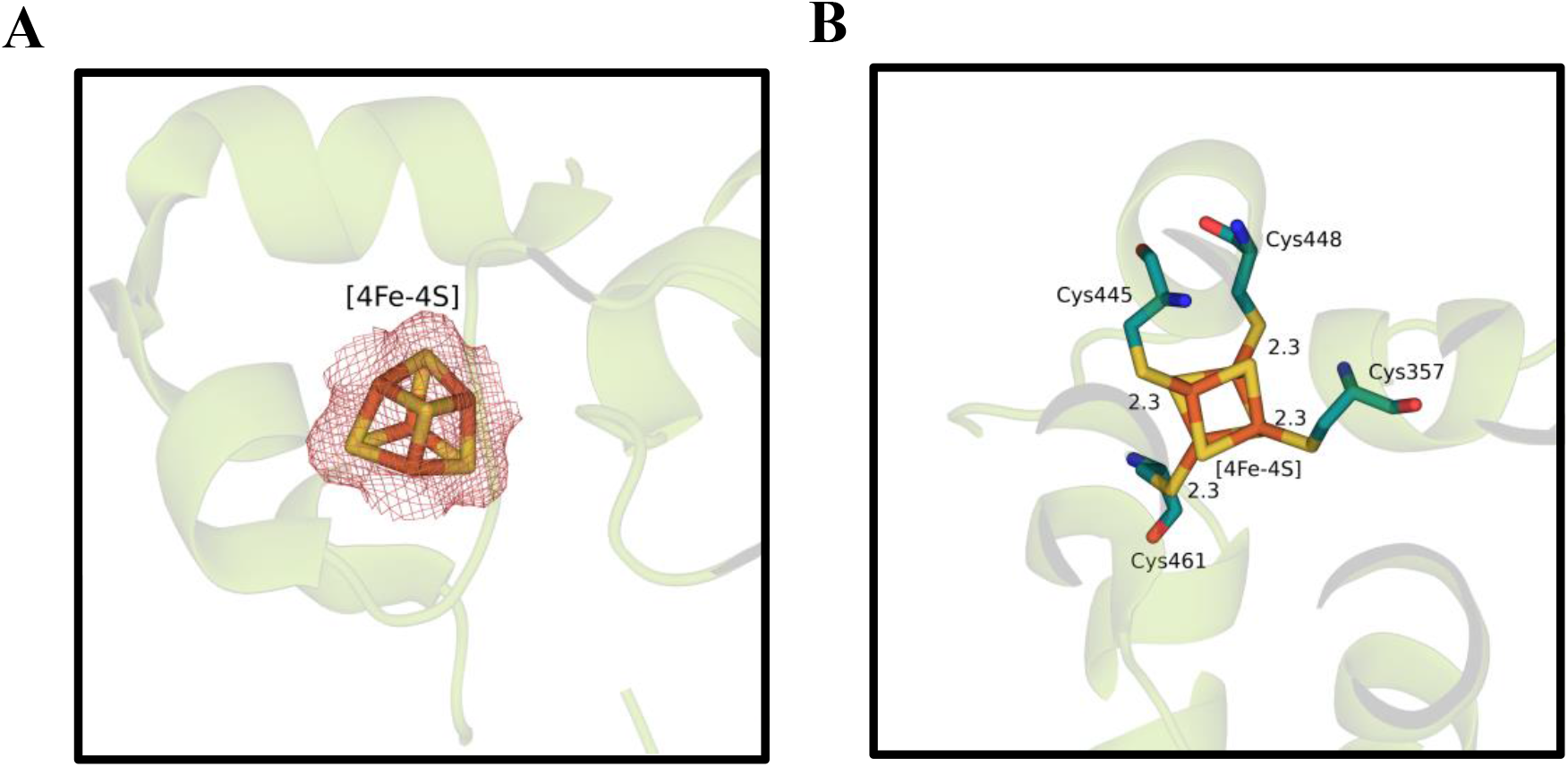
[4Fe-4S] cluster interactions. (A) Electron cloud of [4Fe-4S] cluster at σA-weighted 2 Fo-Fc electron density map contoured at 1.0σ in deepsalmon mesh and *Vc*(6-4) PL cartoon in splitpea color. (B) Direct interactions of [4Fe-4S] clusters with surrounding amino acids of *Vc*(6- 4) PL. Every distance is in units of Angstrom (Å).

[4Fe-4S] clusters are well-known for their oxygen sensitivity, and under aerobic conditions it quickly decomposes into [3Fe-4S] clusters and can further dissociate into [2Fe-2S]. Mostly the decomposition of the cluster deactivates the cluster containing enzymes and handling and purification of these enzymes require using glovebox to prevent decomposition of the oxygen sensitive [4Fe-4S] clusters. In the case of *Vc*(6-4) PL, the entire expression, purification, and crystallization procedures were performed in atmospheric conditions that took months. In our structure, there was no observable damage in the [4Fe-4S] cluster, which might indicate the cluster in the photolyase is unusually oxygen tolerant.

### DNA binding region of *Vc*(6-4) photolyase

The DNA binding region of the *Vc*(6-4) PL has defined electron density covers amino acids Asn178, Phe179, Asp180, Ala181, Asp182, Asn183, Arg184, and Asn185 (**Figure 5A**). In contrast, this region has not a well-defined electron density in *At*(6-4) PL structures obtained in either cryogenic (PDB ID: 4DJA) or ambient temperature (PDB ID: 6DD6) conditions (**Figure 5B** and **C**). A similar analysis were carried out with *Rs*(6-4) PL, where there is a well-defined electron density in the same region (**Figure 5D**). These results suggest that while this region is highly stable in *Rs*(6-4) PL and *Vc*(6-4) PL, it is flexible and disordered in *At*(6-4) PL structure and, therefore, this region has species specific stability differences.

**Figure 5:**
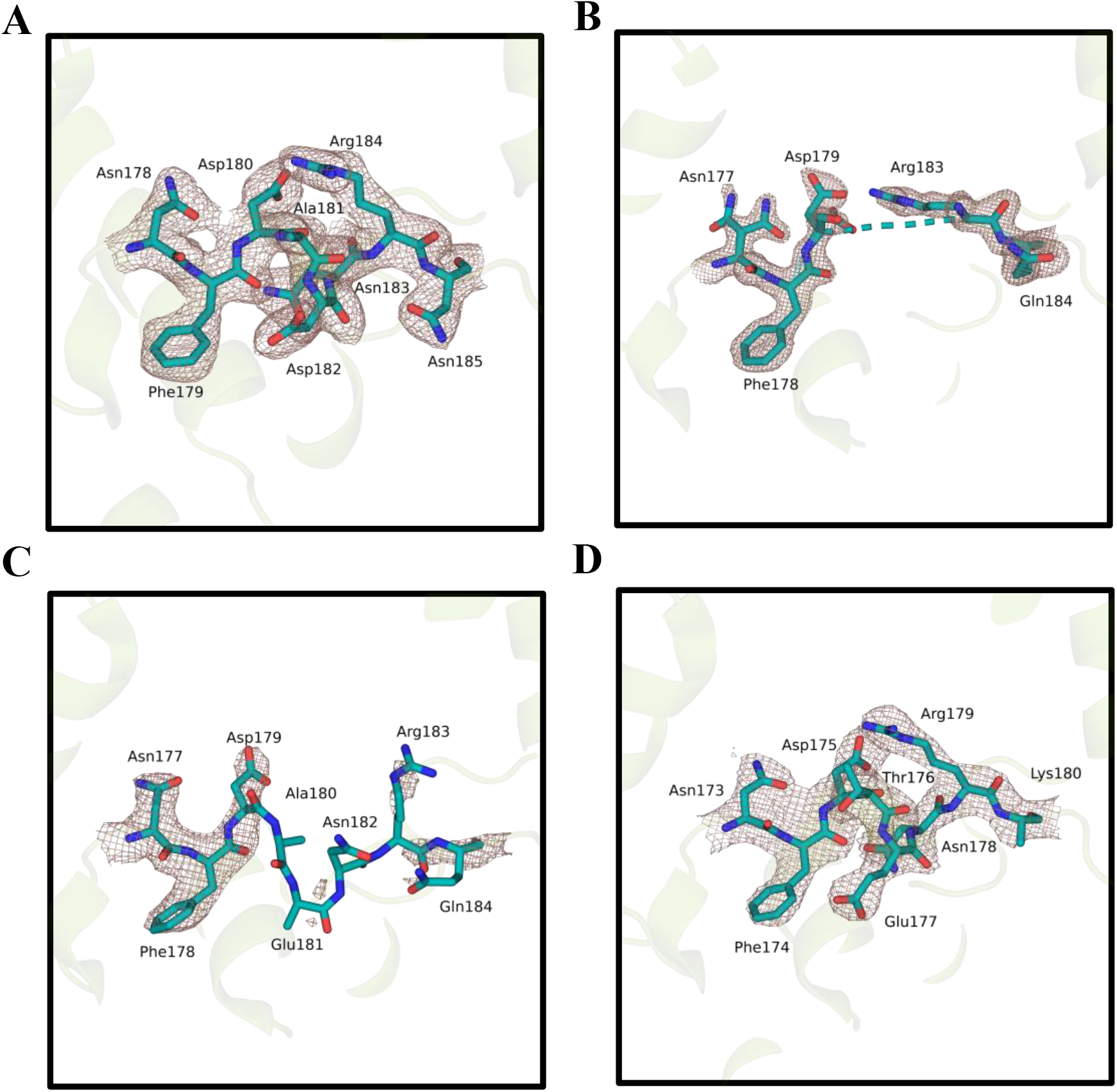
Possible DNA binding site comparison with structurally known wild-type bacterial (6-4) photolyases. Possible DNA binding sites are drawn with electron clouds at σA-weighted 2 Fo-Fc electron density map contoured at 1.0σ in deepsalmon mesh and (6-4) photolyase cartoon in splitpea color. (A) *Vc*(6-4) PL, (B) *At*(6-4) PL cryogenic (PDB ID: 4DJA), (C) *At*(6-4) PL ambient (PDB ID: 6DD6), (D) *Æs*(6-4) PL (PDB ID: 3ZXS). In (B), 3 amino acids are missing, which are Ala180, Glu181, Asn182, due to lack of electron clouds.

Notably, we have not observed a distinct electron density of amino acid residues between 478 and 485 *Vc*(6-4) PL (**Figure 6A**). However, this region has well-defined structures in other bacterial PLs (**Figure 6 B,C and D**). Interestingly, this region is adjacent to the 23^rd^ helix, which is one of the positively surface charge helices around the catalytic region (**Figure 7A**). When we superimposed *Vc*(6-4) PL with *Dm*(6-4) PL (PDB ID: 3CVU) with (6-4) T-T, this region and 23^rd^ helix were seen to pass through damaged DNA (**Figure 7B**). This observation suggests that the region between 478 and 485 *Vc*(6-4) PL might be a DNA binding region and further stabilized after interacting with the DNA lesion. Upon investigation of C-terminus of *Vc*(6-4) PL and *Dm*(6-4) PL, 2 additional helices were identified on *Vc*(6-4) PL and the 23^rd^ helix seems to be blocking the DNA lesion in superimposed structure. Further analysis of superimposed structures indicated that the catalytic cofactor FADs are in similar conformation and their RMSD is 0.7 Å (**Figure 7C**).

**Figure 6:**
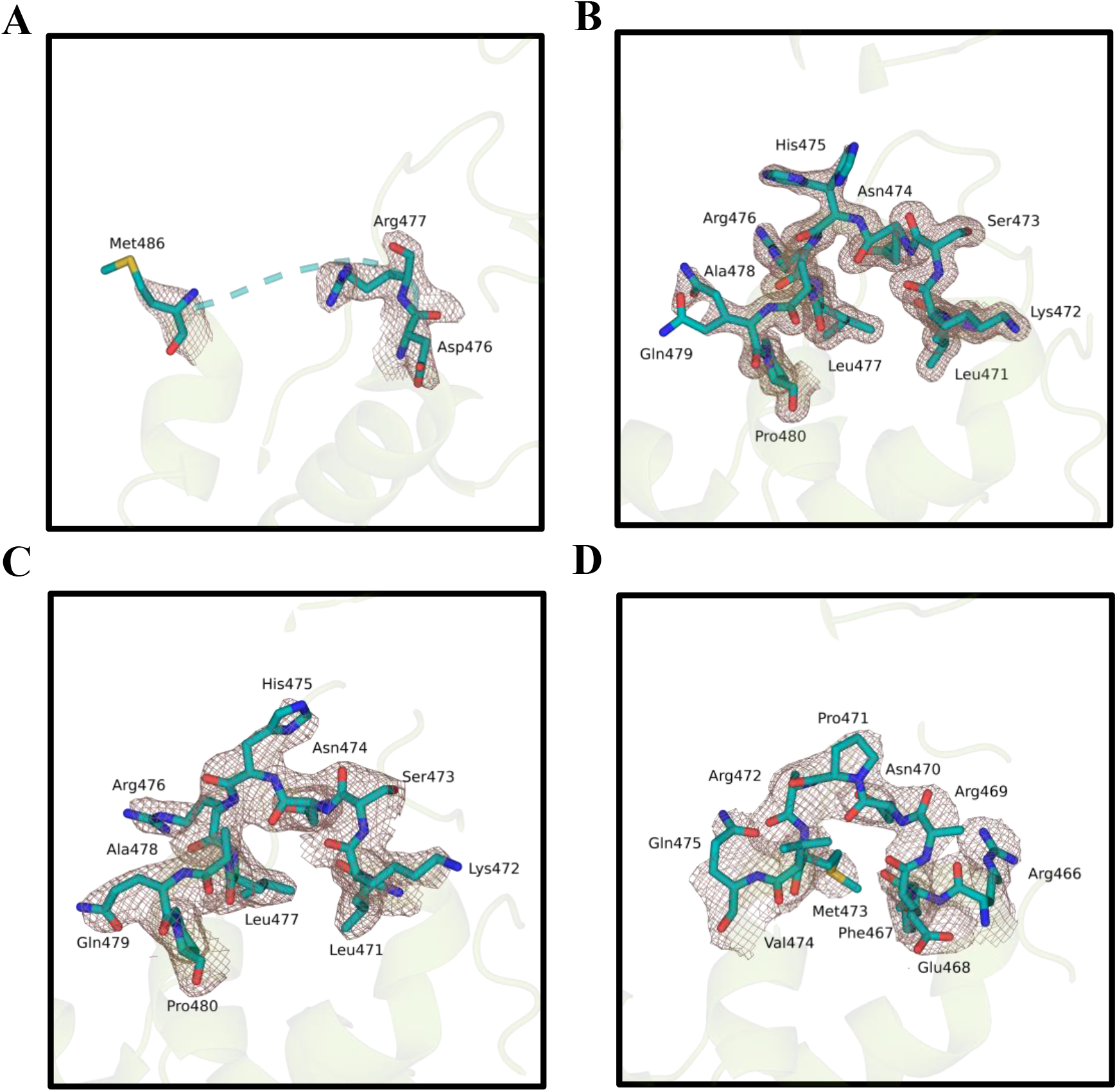
Possible DNA binding domain comparison with structurally known wild-type bacterial (6-4) photolyase. Each possible DNA binding site’s electron cloud is drawn at σA- weighted 2 Fo-Fc electron density map contoured at 1.0σ in deepsalmon mesh and (6-4) photolyase cartoon in splitpea color. (A) *Vc*(6-4) PL, (B) *At*(6-4) PL cryogenic (PDB ID: 4DJA), (C) *At*(6-4) PL ambient (PDB ID: 6DD6), (D) *Rs*(6-4) PL (PDB ID: 3ZXS). In (A), 8 amino acids are missing, which are Leu478, Ala479, Asn480, Asn481, Pro482, Arg483, Ile484, Gly485 due to lack of electron density cloud.

**Figure 7:**
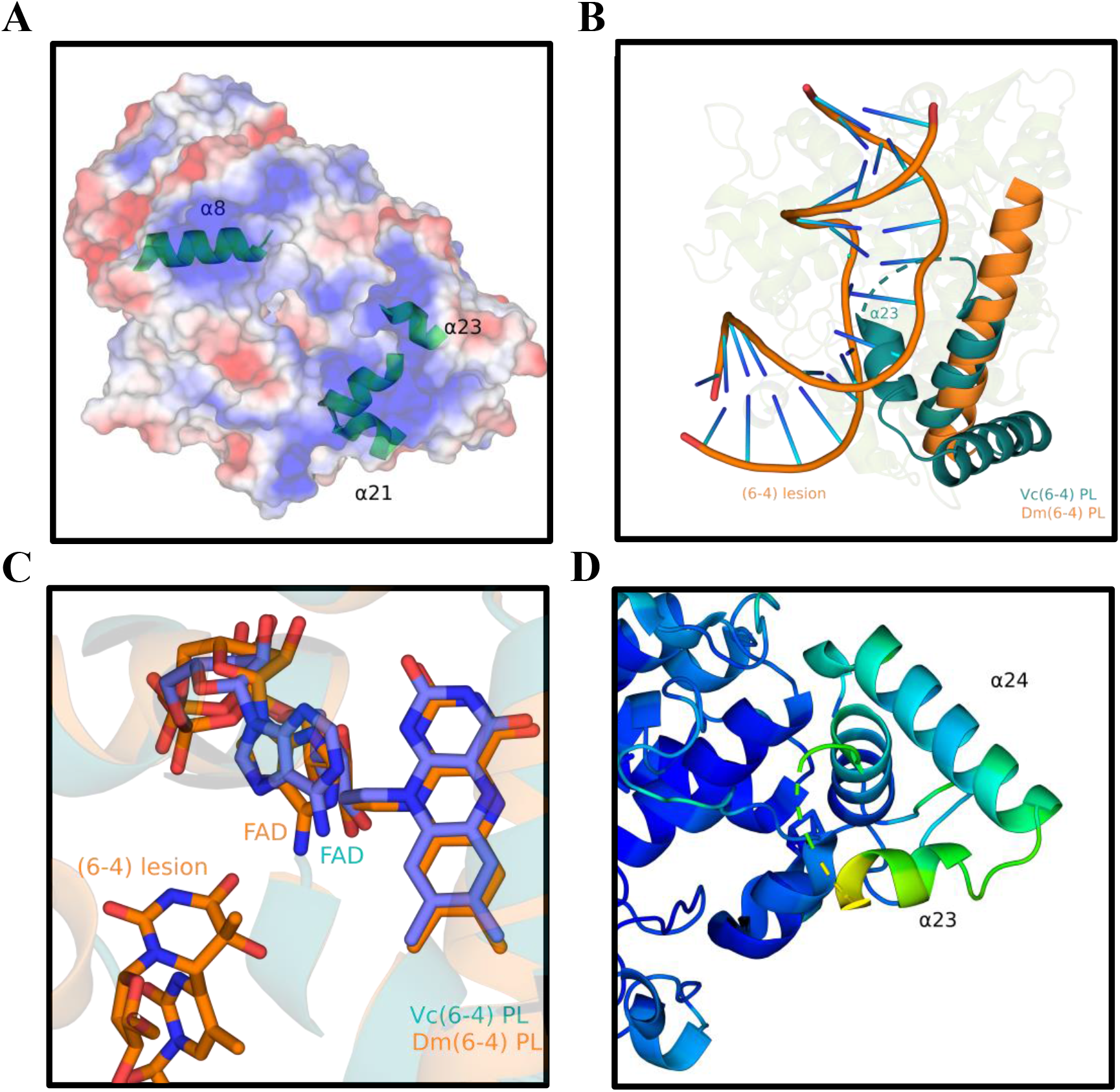
DNA binding comparison between *Vc*(6-4) PL and *Dm*(6-4) PL. (A) Generated surface density charge of *Vc*(6-4) PL, where blue is positive while red is negatively charged. Helices with positive surface charge are indicated in the structure. (B) Superimposed (RMSD: 3.944 Å) image of *Vc*(6-4) PL and *Dm*(6-4) PL with (6-4) DNA lesion (PDB ID: 3CVU). Deepteal is for *Vc*(6-4) PL, while Tv_orange is for *Dm*(6-4) PL. (C) FAD comparison with superimposed images of *Vc*(6-4) PL and *Dm*(6-4) PL. All atoms RMSD between the structures for FAD cofactor is 0.7 Å. (D) B-factor analysis of the C-terminal of *Vc*(6-4) PL, where the highest b-factor has a yellowish color, while the lowest b-factor has a darker blue color. α23 has the highest b-factor in the structure.

To DNA lesions to bind the bacterial (6-4) PL, while still interacting with the FAD, there may be conformational changes on the 23^rd^ helix to provide a gap for DNA lesion. B-factor analysis showed that, 23^rd^ helix has the highest b-factor values in the structure and therefore, it has the highest probability to subject a conformational change (**Figure 7D**). According to the superimposition of DNA lesion from *Dm*(6-4) PL with *Vc*(6-4) PL, which is compatible with a prototypical photolyase-DNA binding mode, helix α23 of *Vc*(6-4) PL interacts with the DNA, while helix α24 looks away from the DNA lesion. The surface charge of helix α23 is positive, which is consistent with the role in DNA binding. The last two helixes (α23, α24) are missing in *Dm*(6-4) PL or most of the other PLs, which indicates they may play a regulatory role rather than their involvement in repair activity.

## Discussion

Photolyases (PLs) belong to a large family of Cryptochrome/Photolyase responsible for repairing the UV-induced DNA damages. Different types of photolyases have been discovered and they are shown to repair two different types of DNA damages: cyclobutane pyrimidine dimers (CPD) (Pyr<>Pyr) and pyrimidine-pyrimidone photoproducts (Pyr[6–4]Pyr). CPD photoproducts are mainly repaired by CPD PLs while (6-4) photoproducts are being repaired by (6-4) PLs (12). The reaction mechanism of CPD photolyases is elucidated at *in vivo* level (1, 5 & 6). On the other hand, different reaction mechanisms are proposed for the (6-4) PL. The (6-4) PLs is initially thought to specifically present in eucaryotes (13). However, studies revealed that bacteria also possess (6-4) PLs (15, 16 & 31). Comparison of the crystal structure of the *Vc* (6-4) PL with other known bacterial (6-4) PLs indicated that cofactor binding regions are well conserved with some differences: a water molecule interacts with the Asp106 in *Vc*(6-4) PL with a very short distance compare to the *At*, and *Rs*(6-4) PLs, where water molecule interacts with Gly105 (**Figure 2B**). The FAD binding region of the *Vc*(6-4) PL is highly conserved. However, one notable difference is that Glu410 makes a hydrogen bond with Asp395; however, in the case of Glu399 (corresponds to Glu410 of *Vc*(6-4) PL) makes a salt bridge with His384 (corresponds to Asp395 of *Vc*(6-4) PL) *At*(6-4) PL and *Rs*(6-4) PL (13, 16). Whether such structural difference affects its activity or conformation of FAD needs to be further investigated. The His-His-X-X-Arg motif in (6-4) PLs is shown to be important for their catalytic activity (16). This motif is also conserved in the *Vc*(6- 4) PL. Further conformational comparison of the *Vc*(6-4) PL and *Dm*(6-4) PL of this motif indicates His373 and Arg376 of *Dm*(6-4) PL have identical configuration while His372 (His364 of *Dm*(6-4) PL) acquired a different configuration, which may reveal similar/different functionalities between species.

One of the major differences between *Vc*(6-4) PL and *At*(6-4) PL structures is their predicted DNA binding site (**Figure 5**). The predicted site (amino acid residues between 178 and 185) of Vc(6-4) PL has a distinct structure compared to *At*(6-4) PL, where the corresponding region has no well-defined structure (17). The same region of *Rs*(6-4) PL has also a well-ordered electron density map except its Arg residue (16), which indicates species specific stability differences of DNA binding domains. Comparison of *Vc*(6-4) PL and *At*(6-4) PL revealed that *Vc*(6-4) PL has an unstructured region with 8 amino acid residues (**Figure 6A**), which might be DNA binding region and also conserved in other bacterial PLs. Arg483 might form a salt bridge with (6-4) photoproduct that stabilizes the disordered region. The general structure of *Vc* and *Dm*(6-4) PLs are highly similar except in C terminus, where *Vc*(6-4) PLs contain additional 2 helixes. Due to the position of the 23^rd^ helix, DNA lesions cannot bind the *Vc*(6-4) PL similar to *Dm*(6-4) PL (**Figure 7B**). However, the FADs are located in an extremely similar manner, which indicates a similar DNA lesion binding between the PLs (**Figure 7C**). A conformational change upon DNA binding at the *Vc*(6-4) PL might open a space for DNA to bind. The 23^rd^ helix is the main candidate for the conformational change as it is the major blocker of the DNA and has the largest b-factor (**Figure 7D**).

### Experimental procedures

#### Expression and purification

Full-length (6-4) photolyase from *Vibrio cholerae* O1 bivar Tor str. N16961 in pET28a(+) vector between NdeI and BamHI cut sites purchased from Genscript (Piscataway, NJ, USA) as codon optimized for *Escherichia coli* protein expression. The plasmid transformed into *E. coli* BL21 Rosetta-2 strain and grown in 4.5 liters of LB-Miller liquid growth media supplemented with 50 μg/mL Kanamycin and 35 μg/mL Chloramphenicol antibiotics at 37 °C with 110 rpm shaking. When OD_600_ reached to 0.8, induction of bacterial protein expression was performed by adding 0.4 mM of isopropyl *β-D*-thiogalactopyranoside (IPTG) as the final concentration, and induction was performed at 18 °C for 24 hours. The culture was centrifuged at 2850 g for 45 minutes at 4 °C and the pellet was stored at −80 °C until further use. Lysis buffer containing 500 mM NaCl, 50 mM Tris-HCl pH 7.5, 10% glycerol (v/v), 0.1% Triton X-100 (v/v) was added into pellets and sonication (Branson W250 sonifier, USA) was performed for 45 seconds at 60% power 3 times. Samples were ultracentrifuged at 35000 rpm with Ti-45 rotor (Beckman, USA) for 1 hour at 4 °C, subsequently the supernatant was filtered with 45 μm cellulose mixed ester filter (ISOLAB, Germany), then applied to Ni-NTA agarose resin (QIAGEN, USA). The column was equilibrated with His A buffer containing 200 mM NaCl, 20 mM Tris-HCl pH 7.5 (v/v), 5% glycerol (v/v) then sample was loaded with 2.5 ml/min flow rate. Washing of the column was performed with His A buffer, and protein was eluted with His B buffer containing 200 mM NaCl, 20 mM Tris-HCl pH 7.5, 250 mM imidazole, 5% glycerol (v/v) into 5 ml of 100% glycerol to prevent protein precipitation. Yellow colored (6-4) photolyase was flash frozen with liquid nitrogen and stored at −80 °C with the final concentration of 25% glycerol (v/v).

#### Crystallization and Harvesting

Initial crystallization screening experiments were setup with 5 mg/ml final concentration of *Vc*(6-4) photolyase with the sitting drop, microbatch screening (under oil) method at 4 °C by mixing 0.83 μl of protein with equal volume of commercial crystal screening conditions in 72 well Terasaki plates. The well containing the protein:cocktail mixture was covered with 16.6 μl of parafilm oil to allow slow evaporation of solvents. Approximately 3500 commercially available crystal screening conditions were tested, and the yellow-colored crystals were observed at Wizard Synergy #40 (Rigaku, Japan), which contains 2 M of ammonium citrate/citric acid pH 7.5 and 5% PEG 400 (v/v) after 7 weeks. The conditions were further optimized by mixing 0.5 μl Cryo-Pro #45 (Hampton research, USA), which contain 1 M sodium sulfate decahydrate, 1 μl of protein, 1 μl of Wizard Synergy #40. The crystals were flash frozen by quickly plunging them into liquid nitrogen, and data collection was performed at 100 K.

#### Data collection and Processing

Two X-ray diffraction datasets were collected to 2.0 Å and 2.5 Å resolutions from two large crystals with a wavelength of 1.54 Å at University of Health Sciences (Istanbul, Turkiye) with Rigaku’s XtaLAB Synergy Flow X-ray diffractometer (Rigaku, Japan). The PhotonJet-R (Rigaku, Japan) X-ray generator was operated at 40 kV and 30 mA with 23% beam intensity to mitigate the radiation damage during data collection. HyPix-Arc-150° detector (Rigaku, Japan) was used with a 60 mm detector distance. To further minimize the exposure time we prefered not to collect fine-sliced ossilation data rather 1-degree oscillation scan width low X-ray dose chosen with 15 and 30 seconds exposure times. The total run time for the two data collections was 1 h 11 m 30 sec, and 2 h 33 min 0 sec respectively. Profit merge is done by using the data from both crystals and data reduction is performed with CrysAlis Pro (32) software version 171.42.51a. Two datasets were merged with 99.6 % completeness and 60.6 fold multiplicity. Unit cell dimensions were a=200.8 Å b=200.8 Å c=77.0 Å α=90 β=90 γ=120, in space group P64 22.

#### Structure Determination and Refinement

Swiss model (33) was used for building a search model for structure determination that gave the best result with PDB ID: 4DJA (13), which was used in the automated molecular replacement program *PHASER* (34) in *PHENIX* (35) software version 1.20.1-4487. Further refinements were performed with *PHENIX*, and the addition of water and cofactors was performed by using *COOT* (36) software version 0.9.6. The final R_work_ is 22.54% and R_free_ is 28.30%, completeness is 99.60% with 24.250 to 2.500 Å refinement resolution (Table 1). The structure contains no Ramachandran outliers with 98.01% of the residues in the favored regions. The structure was deposited to the RSCB PDB website with the PDB ID: 7YKN. Figures were generated at *PyMOL* (37) Version 2.4.1. Data collection and structure determination information are summarized in Table 1.

**Table 1:** Statistics for data collection, processing and structure refinement.

|  |  |
| --- | --- |
| Beamline | Turkish Light Source (*Turkish DeLight*) |
| Resolution range | 24.25-2.5 (2.589-2.5) |
| Space group | P6<sub>4</sub> 2 2 |
| Unit cell | 200.8, 200.8, 77.0, 90, 90, 120 |
| Total reflections | 1935882 (189833) |
| Unique reflections | 32043 (3130) |
| Multiplicity | 60.4 (60.6) |
| Completeness (%) | 99.60 (99.97) |
| Mean I/$\sigma$ | 5.61 (1.70) |
| Wilson B-factor | 15.97 |
| R-merge | 1.382 (3.765) |
| R-meas | 1.394 (3.796) |
| R-pim | 0.1776 (0.4855) |
| CC 1/2 | 0.955 (0.546) |
| CC* | 0.988 (0.84) |
| Reflections used in refinement | 31969 (3130) |
| Reflection used for R-free | 1068 (105) |
| R-work | 0.2254 (0.2891) |
| R-free | 0.2830 (0.3620) |
| CC(work) | 0.910 (0.712) |
| CC(free) | 0.929 (0.542) |
| Number of non-hydrogen atoms | 4778 |
| Macromolecules | 4120 |
| Ligands | 101 |
| solvent | 557 |
| Protein residues | 507 |
| RMS(bonds) | 0.002 |
| RMS(angles) | 0.48 |
| Ramachandran favored (%) | 98.01 |
| Ramachandran allowed (%) | 1.99 |
| Ramachandran outliers (%) | 0.00 |
| Rotamer outliers (%) | 1.84 |
| Clashscore | 5.92 |
| Average B-factor | 22.46 |
| macromolecules | 22.31 |
| Ligands | 17.61 |
| solvent | 24.49 |
| Number of TLS groups | 8 |

## Supporting information

Response Letter

## Data Availability

All data generated or analyzed during this study are included in this article and the structure was deposited to the RCSB Protein Data Bank website with the PDB ID: 7YKN.

## Author Contributions

H.D. and I.H.K. designed and coordinated the project. H.D guided B.C. to execute all the sample preparation, data collection, data reduction and refinement of photolyase structure. H.D., I.H.K and B.C analyzed data. H.D., I.H.K and B.C. prepared the manuscript.

## Acknowledgements

Authors would like to dedicate this manuscript to the memory of Dr. Albert E. Dahlberg and Dr. Nizar Turker. H.D. acknowledges support from NSF Science and Technology Center grant NSF- 1231306 (Biology with X-ray Lasers, BioXFEL). The authors gratefully acknowledge use of the services and Turkish Light Source (*Turkish DeLight*) X-ray facility at Sağlık Bilimleri University Deneysel Tip Araştirma ve Uygulama Merkezi (SBU-DETUAM). The authors also gratefully acknowledge use of the services and facilities at Koç University Isbank Research Centre for Infectious Diseases (KUIS-CID). This publication has been produced benefiting from the 2232 International Fellowship for Outstanding Researchers Program of the TÜBİTAK (Project No. 118C270). However, the entire responsibility of the publication belongs to the authors of the publication. The financial support received from TÜBİTAK does not mean that the content of the publication is approved in a scientific sense by TÜBİTAK. IHK would like to thank the Istanbul Development Agency grant (ISTKA-TR/14/EVK/0039) for their financial support.

---

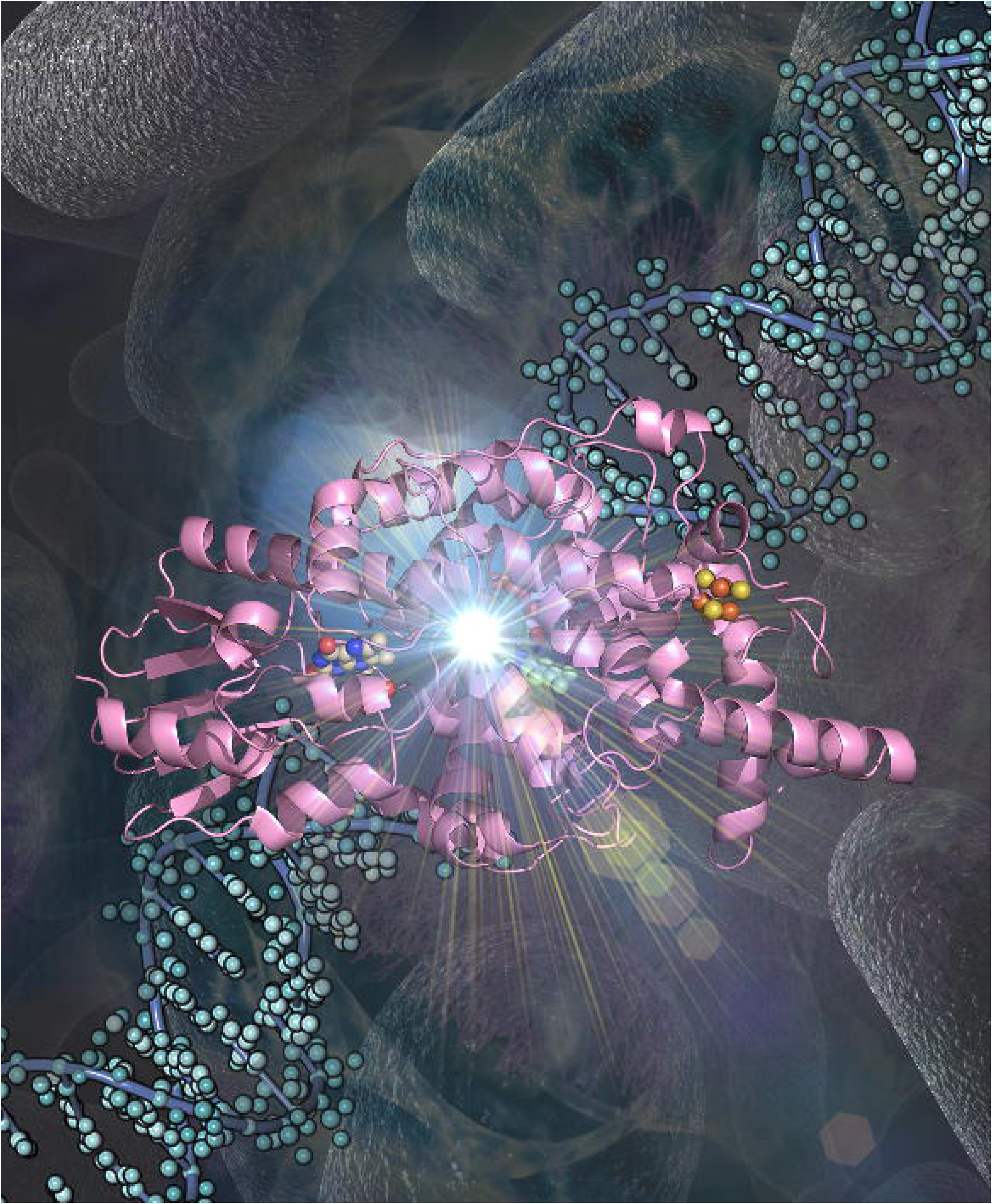

## References

1. Kavakli, I. H., Baris, I., Tardu, M., Gul, S., Oner, H., Cal, S., Bulut, S., Yarparvar, D., Berkel, C., Ustaoglu, P., and Aydin, C. (2017) The Photolyase/Cryptochrome Family of Proteins as DNA Repair Enzymes and Transcriptional Repressors. Photochem Photobiol 93, 93–103

2. Sancar, A. (2004) Photolyase and cryptochrome blue-light photoreceptors. Adv Protein Chem 69, 73–100

3. Chaves, I., Pokorny, R., Byrdin, M., Hoang, N., Ritz, T., Brettel, K., Essen, L. O., van der Horst, G. T., Batschauer, A., and Ahmad, M. (2011) The cryptochromes: blue light photoreceptors in plants and animals. Annu Rev Plant Biol 62, 335–364

4. Kavakli, I. H., and Sancar, A. (2004) Analysis of the role of intraprotein electron transfer in photoreactivation by DNA photolyase in vivo. Biochemistry 43, 15103–15110

5. Kavakli, I. H., Ozturk, N., and Gul, S. (2019) DNA repair by photolyases. Adv Protein Chem Struct Biol 115, 1–19

6. Sancar, A. (2016) Mechanisms of DNA Repair by Photolyase and Excision Nuclease (Nobel Lecture). Angew Chem Int Ed Engl 55, 8502–8527

7. Zhong, D. (2015) Electron transfer mechanisms of DNA repair by photolyase. Annu Rev Phys Chem 66, 691–715

8. Asimgil, H., and Kavakli, I. H. (2012) Purification and characterization of five members of photolyase/cryptochrome family from Cyanidioschyzon merolae. Plant Sci 185-186, 190–198

9. Ozturk, N. (2017) Phylogenetic and Functional Classification of the Photolyase/Cryptochrome Family. Photochem Photobiol 93, 104–111

10. Ozturk, N., Kao, Y. T., Selby, C. P., Kavakli, I. H., Partch, C. L., Zhong, D., and Sancar, A. (2008) Purification and characterization of a type III photolyase from Caulobacter crescentus. Biochemistry 47, 10255–10261

11. Selby, C. P., and Sancar, A. (2006) A cryptochrome/photolyase class of enzymes with singlestranded DNA-specific photolyase activity. Proc Natl Acad Sci U S A 103, 17696–17700

12. Sancar, A. (2003) Structure and function of DNA photolyase and cryptochrome blue-light photoreceptors. Chem Rev 103, 2203–2237

13. Zhang, F., Scheerer, P., Oberpichler, I., Lamparter, T., and Krauss, N. (2013) Crystal structure of a prokaryotic (6-4) photolyase with an Fe-S cluster and a 6,7-dimethyl-8-ribityllumazine antenna chromophore. Proc Natl Acad Sci US A 110, 7217–7222

14. Zhang, M., Wang, L., and Zhong, D. (2017) Photolyase: Dynamics and electron-transfer mechanisms of DNA repair. Arch Biochem Biophys 632, 158–174

15. Dikbas, U. M., Tardu, M., Canturk, A., Gul, S., Ozcelik, G., Baris, I., Ozturk, N., and Kavakli, I. H. (2019) Identification and Characterization of a New Class of (6-4) Photolyase from Vibrio cholerae. Biochemistry 58, 4352–4360

16. Geisselbrecht, Y., Fruhwirth, S., Schroeder, C., Pierik, A. J., Klug, G., and Essen, L. O. (2012) CryB from Rhodobacter sphaeroides: a unique class of cryptochromes with new cofactors. EMBO Rep 13, 223–229

17. Oberpichler, I., Pierik, A. J., Wesslowski, J., Pokorny, R., Rosen, R., Vugman, M., Zhang, F., Neubauer, O., Ron, E. Z., Batschauer, A., and Lamparter, T. (2011) A photolyase-like protein from Agrobacterium tumefaciens with an iron-sulfur cluster. PLoS One 6, e26775

18. Park, H. W., Kim, S. T., Sancar, A., and Deisenhofer, J. (1995) Crystal structure of DNA photolyase from Escherichia coli. Science 268, 1866–1872

19. Tamada, T., Kitadokoro, K., Higuchi, Y., Inaka, K., Yasui, A., de Ruiter, P. E., Eker, A. P., and Miki, K. (1997) Crystal structure of DNA photolyase from Anacystis nidulans. Nat Struct Biol 4, 887–891

20. Komori, H., Masui, R., Kuramitsu, S., Yokoyama, S., Shibata, T., Inoue, Y., and Miki, K. (2001) Crystal structure of thermostable DNA photolyase: pyrimidine-dimer recognition mechanism. Proc Natl Acad Sci U S A 98, 13560–13565

21. Hitomi, K., DiTacchio, L., Arvai, A. S., Yamamoto, J., Kim, S. T., Todo, T., Tainer, J. A., Iwai, S., Panda, S., and Getzoff, E. D. (2009) Functional motifs in the (6-4) photolyase crystal structure make a comparative framework for DNA repair photolyases and clock cryptochromes. P Natl Acad Sci USA 106, 6962–6967

22. Worthington, E. N., Kavakli, I. H., Berrocal-Tito, G., Bondo, B. E., and Sancar, A. (2003) Purification and characterization of three members of the photolyase/cryptochrome family bluelight photoreceptors from Vibrio cholerae. J Biol Chem 278, 39143–39154

23. Tardu, M., Bulut, S., and Kavakli, I. H. (2017) MerR and ChrR mediate blue light induced photooxidative stress response at the transcriptional level in Vibrio cholerae. Sci Rep 7, 40817

24. Atalay, N., Akcan, E. K., Gül, M., Ayan, E., Destan, E., Ertem, F. B., Tokay, N., Çakilkaya, B., Nergiz, Z., Karakadioğlu, G., Kepceoğlu, A., Yapici, İ., Tosun, B., Baldir, N., Yildirim, G., Johnson, J. A., Güven, Ö., Shafiei, A., Arslan, N. E., Yilmaz, M., Kulakman, C., Paydos, S. S., Çinal, Z. S., Şabanoğlu, K., Pazarçeviren, A., Yilmaz, A., Canbay, B., Aşci, B., Kartal, E., Tavli, S., Çaliseki, M., Göç, G., Mermer, A., Yeşilay, G., Altuntaş, S., Tateishi, H., Otsuka, M., Fujita, M., Tekin, Ş., Çiftçi, H., Durdaği, S., Dinler Doğanay, G., Karaca, E., Kaplan Türköz, B., Kabasakal, B. V., Kati, A., & Demirci, H. (2022) Cryogenic X-ray crystallographic studies of biomacromolecules at Turkish Light Source “Turkish DeLight”. Preprint at https://doi.org/10.1101/2022.09.03.506456.

25. Gul, M., Ayan, E., Destan, E., Johnson, J. A., Shafiei, A., Kepceoğlu, A., Yilmaz, M., Ertem, F. B., Yapici, I., Tosun, B., Baldir, N., Tokay, N., Nergiz, Z., Karakadioğlu, G., Paydos, S. S., Kulakman, C., Ferah, C. K., Güven, Ö., Atalay, N., … DeMirci, H. (2022) Rapid and high resolution ambient temperature structure determination at Turkish light source. Preprint at https://doi.org/10.1101/2022.10.12.511637

26. Faraji, S., and Dreuw, A. (2017) Insights into Light-driven DNA Repair by Photolyases: Challenges and Opportunities for Electronic Structure Theory. Photochemistry and Photobiology 93, 37–50

27. Matsubara, T., Araida, N., Hayashi, D., and Yamada, H. (2014) Computational Study on the Mechanism of the Electron-Transfer-Induced Repair of the (6-4) T-T Photoproduct of DNA by Photolyase: Possibility of a Radical Cation Pathway. B Chem Soc Jpn 87, 390–399

28. Maul, M. J., Barends, T. R. M., Glas, A. F., Cryle, M. J., Domratcheva, T., Schneider, S., Schlichting, I., and Carell, T. (2008) Crystal Structure and Mechanism of a DNA (6-4) Photolyase. Angew Chem Int Edit 47, 10076–10080

29. Hitomi, K., Nakamura, H., Kim, S. T., Mizukoshi, T., Ishikawa, T., Iwai, S., and Todo, T. (2001) Role of two histidines in the (6-4) photolyase reaction. Journal of Biological Chemistry 276, 10103–10109

30. Li, J., Liu, Z., Tan, C., Guo, X., Wang, L., Sancar, A., and Zhong, D. (2010) Dynamics and mechanism of repair of ultraviolet-induced (6-4) photoproduct by photolyase. Nature 466, 887–890

31. Hendrischk, A. K., Fruhwirth, S. W., Moldt, J., Pokorny, R., Metz, S., Kaiser, G., Jager, A., Batschauer, A., and Klug, G. (2009) A cryptochrome-like protein is involved in the regulation of photosynthesis genes in Rhodobacter sphaeroides. Mol Microbiol 74, 990–1003

32. Diffraction, R. O. (2021) CrysAlisPro Software system, version 1.171.42.51a, Rigaku Corporation, Wroclaw, Poland.

33. Bienert, S., Waterhouse, A., de Beer, T. A., Tauriello, G., Studer, G., Bordoli, L., and Schwede, T. (2017) The SWISS-MODEL Repository-new features and functionality. Nucleic Acids Res 45, D313–D319

34. Mccoy, A. J., Grosse-Kunstleve, R. W., Adams, P. D., Winn, M. D., Storoni, L. C., and Read, R. J. (2007) Phaser crystallographic software. J Appl Crystallogr 40, 658–674

35. Adams, P. D., Afonine, P. V., Bunkoczi, G., Chen, V. B., Davis, I. W., Echols, N., Headd, J. J., Hung, L. W., Kapral, G. J., Grosse-Kunstleve, R. W., McCoy, A. J., Moriarty, N. W., Oeffner, R., Read, R. J., Richardson, D. C., Richardson, J. S., Terwilliger, T. C., and Zwart, P. H. (2010) PHENIX: a comprehensive Python-based system for macromolecular structure solution. Acta Crystallogr D Biol Crystallogr 66, 213–221

36. Emsley, P., and Cowtan, K. (2004) Coot: model-building tools for molecular graphics. Acta Crystallogr D Biol Crystallogr 60, 2126–2132

37. DeLano, W. L. (2009) PyMOL molecular viewer: Updates and refinements. Abstr Pap Am Chem S 238

