## Supplementary material for "The crystal structure of *Vibrio cholerae* (6-4) photolyase reveals interactions with cofactors and a DNA binding region": Response Letter

**Point by Point Responses to Reviewers' comments**

**Please find below our point-by-point responses to our Reviewers' comments written in blue text. All changes on the manuscript are also highlighted with yellow color on the main text and supplementary files.**

**Manuscript Number: [redacted]**
**Crystal structure of Vibrio cholerae (6-4) photolyase with DNA binding region**

**Significance**

Reviewer 1: This is an outstanding manuscript that describes the crystal structure of a (6-4) photolyase from an unexpected source.It greatly contributes to the understanding of this class of enzymes.

**RESPONSE: We sincerely would like to express our gratitude for the kind words of our Reviewer 1.**

**Critique**

Reviewer 1: The authors purify and crystallize a (6-4) photolyase from Vibrio cholera, solve the structure, identify the chromophore and propose a plausible mechanism of reaction. Extremely well-done.

**RESPONSE: We again would like to thank our Reviewer 1 for his/her encouraging comments on the manuscript.**

I suggest the authors provide a structure figure for [redacted] cover.

**RESPONSE: As suggested by our Reviewer 1 we prepared a [redacted] cover image and submitted for its consideration by the [redacted] editorial board.**

Please provide an overall rating of this study as it currently stands.

Reviewer 1: Exceptional and of broad interest

**Significance**

Reviewer 2: This review is focused on the crystallographic analysis performed in this study.

**Critique**

Reviewer 2: There appear to be no major problems with the crystallography. The data set was collected with very high redundancy/multiplicity (60 fold), and this seems to have driven the R-merge statistic to a very high value of 138.3% (over all data). This makes it difficult to know whether there are any issues with the Space Group assignment. It would be useful to know what the R-merge value was for the lowest resolution shell. And perhaps not all of the data needs to be used? Was there any sign of radiation damage? Perhaps the iron sulfur cluster is more sensitive to damage, and could benefit from fewer images being included (say a multiplicity of ~6 or so)? Given that the Crystallographic refinement R factors are reasonable, and the maps look fine, there probably is not a problem with the Space Group assignment, but it would be useful to know more about the data collection strategy, and whether the suggested considerations might improve the maps/model.

**RESPONSE: We would like to thank our reviewer for his/her valuable comments on redundancy and R-merge. We apologize for not being able to explain our story line clearly in the main text. Obtaining well diffracting Vc(6-4) PL crystals were extremely difficult and after exhaustively screening 3500 sparse matrix and grid screen conditions only one condition yielded crystals. We further optimized this condition for further improving diffraction quality however no single crystal was enough to yield a complete dataset. We merged two 360 degrees datasets to obtain the final merged dataset that we used for further refinement. We shared the same Radiation damage concern of our reviewer and we attenuated the beam to %23 to mitigate the radiation damage. The data collection strategy was optimized by CrysAlisPro software installed at the Turkish DeLight beamline. We attempted to merge less image with lower multiplicity, however, the best electron density maps with optimum Rmerge and crystallographic refinement R factors were obtained when we include all collected data. We used a home X-ray source; a better low redundant dataset collection strategy could have been employed at a syncrothron facility. At the moment we do not have a national or international carrier that can transport our cryo dewars to international locations and this limited our access and data collection at a synchrotron facility.**

**Following the advice of our Reviewer 2, we redid the data processing and reduction part by using one sixth of the reflections from one of the data sets to reduce multiplicity. As the reviewer predict, we lowered the Rmerge to 0.988 (2.506) when multiplicity is 10.2 (10.1). However, we did not observe a change in space group suggestions on CrysAlisPro software. We also lost some completeness %98.77 (91.60), while increasing Rwork to 0.2371 (0.4303) and Rfree to 0.2802 (0.4428). We did not observe any drastic changes in the crystal structure and electron density of cofactors but we lost the electron density of some positively charged side chains in the DNA binding region, which may further support our point that the enzyme is extremely flexible. When multiplicity is reduced, we are sacrificing some critical information in the highest electron shell (2.5-2.59 Å) therefore, our best option was to merge two different crystal data sets to increase the resolution of our data. If we reduce the resolution to 3.5 Å with one twentieth of the reflections then Rmerge reduces to 0.61 (1.724) with 9.6 (10.2) multiplicity. We lose crucial information on side chain conformations and almost all the water molecule electron clouds. By considering these, the option of using high multiplicity for the sake of higher resolution was the best option for us.**

**To reflect our reviewer's concern we completely re-written the last paragraph of the "Crystallization, Data collection and Processing" text as follows:**

**"Data collection and Processing**

**Two X-ray diffraction datasets were collected to 2.0 Å and 2.5 Å resolutions from two large crystals with a wavelength of 1.54 Å at University of Health Sciences (Istanbul, Turkiye) with Rigaku's XtaLAB Synergy Flow X-ray diffractometer (Rigaku, Japan). The PhotonJet-R (Rigaku, Japan) X-ray generator was operated at 40 kV and 30 mA with 23% beam intensity to mitigate the radiation damage during data collection. HyPix-Arc-150° detector (Rigaku, Japan) was used with a 60 mm detector distance. To further minimize the exposure time we prefered not to collect fine-sliced ossilation data rather, 1-degree oscillation scan width low X-ray dose chosen, with 15 and 30 seconds exposure times. The total run time for the two data collections was 1h 11m 30sec, and 2h 33min 0sec respectively. Profit merge is done by using the data from both crystals and data reduction is performed with CrysAlis Pro (320) software version 171.42.51a. Two datasets were merged with 99.6 % completeness and 60.6 fold multiplicity. Unit cell dimensions were a=200.8 Å b=200.8 Å c=77.0 Å α=90 β=90 γ=120, in space group $P6_4 22$."**

Other points:

Methods, "The structure contains 0.00% Ramachandran outliers with 98.01% favored residues."

Since there are likely to be no outliers, I would change this statistic to read 'The structure contains no Ramachandran outliers with....'

**RESPONSE: As suggested by our Reviewer, this sentence has been changed to:**

***"The structure contains no Ramachandran outliers with 98.01% favored residues."***

Maps are listed in the legends as "σA-weighted 2 F0-Fc" but it should be Fo-Fc (letter o instead of number 0)

**RESPONSE: We sincerely apologize for this typo. We extensively reviewed our manuscript and replaced all the "F0-Fc" with "Fo-Fc" as pointed out by our Reviewer 2.**

Please provide an overall rating of this study as it currently stands.

Reviewer 2: High quality and of interest to a broad readership

**RESPONSE: We would like to thank our Reviewer 2 for considering our manuscript for being high quality and of interest to the broader readership of the [redacted].**
